# Transcriptomic landscape reveals immunity related trade-offs in an invertebrate-fungal host-parasite system

**DOI:** 10.1101/2025.03.28.645961

**Authors:** Paraskevopoulou Sofia, Gattis Sabrina, Ben-Ami Frida

## Abstract

1. By exploiting host resources, parasites impose significant fitness costs onto their hosts, thereby affecting their population dynamics. Hosts, in turn, employ a series of mechanism to resist or tolerate parasitic infections. Unlike vertebrates, which possess a sophisticated immune system, invertebrates rely solely on innate immunity to combat pathogens. Despite over 50 years of research, the molecular basis of innate immunity in non-insect invertebrates remains limited.
2. We used the *Daphnia magna* - *Metschnikowia bicuspidata* host-parasite system to shed light on conserved immune responses among *Daphnia* species and parasite-driven immunological shifts. We examined the transcriptomic landscape across infected, exposed-uninfected and unexposed individuals and identified candidate genes that might be involved in the haemocyte recruitment. Additionally, we identified genes that might encode for reinforcement of the gut epithelium, and thus confer resistance to the parasite.
3. We measured life-history traits and observed that shifts correlated with immune activation. Specifically, exposed-uninfected individuals exhibited a delay in reproductive maturation, likely a direct effect of immune activation. However, these shifts appeared temporal, as animals compensated in total reproductive output over time.
4. Unlike exposed-uninfected animals, infected individuals exhibited metabolic shifts that are indicative of host metabolic exhaustion. This metabolic reallocation aligns with the terminal investment hypothesis, where hosts facing high mortality risk divert resources from somatic maintenance to immediate reproduction in order to maximize fitness before death.
5. Our findings provide novel insights into the molecular and physiological mechanisms underlying invertebrate immune responses and life-history trade-offs in the context of parasitic infections.

## Introduction

Parasites (used here interchangeably with pathogens) impose significant fitness costs to their hosts by exploiting host resources and reducing host growth, reproduction and survival (Marzal et al., 2004; Hall et al., 2009; Pigeault et al., 2018; Eleftheriou, 2020), thereby affecting host population dynamics. Hosts, in turn, employ a series of mechanism to resist or tolerate parasitic infections. Unlike vertebrates, whose immune system is adaptive, invertebrates rely solely on their innate immune system to combat infections (Litman & Cooper, 2007; Litman et al., 2010). This immune system consists of a diverse set of cellular components, such as receptors that recognize pathogen associated molecular patterns (PAMPs), regulators that modulate signaling pathways, and effectors that are directly engaged in the inhibition of pathogen proliferation and survival (Coates et al., 2022; Schlenke & Begun, 2004; Schmid-Hempel, 2009; Rathinam et al., 2024). Yet, the molecular mechanisms central to these defenses remain relatively unexplored in non-insect invertebrates.

The *Daphnia*-microparasites system is ideal for investigating the innate immune response in invertebrates. First, natural populations of *Daphnia* are often infected by a variety of parasites, such as bacterial, fungal, microsporidian and viral pathogens (Ebert, 2005; Goren & Ben-Ami, 2013; Ebert, 2022). Additionally, *Daphnia* species play a crucial role in freshwater ecosystems as keystone species in aquatic trophic webs, and in nutrient cycling through decomposition of organic matter (Ebert, 2022; Gorzelnik et al., 2023). Lastly, *Daphnia* are essential in aquaculture, since they are used as food source for many fish larvae (Das et al., 2012; Miura et al., 2021). Given their ecological and economic importance, parasite infections in *Daphnia* can have cascading effects, disrupting food web stability, altering ecosystem dynamics, and potentially impacting aquaculture industries (Ebert et al., 1997; Decaestecker et al., 2005; Hall et al., 2010; Hall et al., 2011).

Over the past decades, substantial advancements have been made in understanding the ecological and evolutionary impacts of infections in *Daphnia.* Recent studies suggest that infection outcome and parasite virulence are species- and genotype-dependent as well as host age-dependent (Duneau et al., 2011; Hall & Ebert, 2012; Izhar et al., 2015; Ben-Ami, 2019; Izhar et al., 2020; Shaw et al., 2021). In cases of coinfections, the sequence of parasite exposure (i.e., priority effect) has also been found to alter the outcome of these interactions (Ben-Ami et al., 2011; Clay et al., 2019; Manzi et al., 2021; Gurung et al., 2024). Despite these achievements, the molecular mechanisms underlying *Daphnia* immunity remain fairly unexplored, with most molecular studies focusing on infections caused by the bacterium *Pasteuria ramosa*. More precisely, candidate immune genes (e.g., Phenoloxidase) have been identified in the *Daphnia* genome (Labbé & Little, 2009). Later on, McTaggart et al. (2015) identified putative immune genes related to the challenge of *Daphnia magna* hosts with infective and non-infective strains of *P. ramosa*. Recently, it has been shown that distinct resistance loci control infection routes in *Pasteuria*, revealing a complex genetic architecture that underlines host-parasite interactions (Bégné et al., 2024). Further transcriptomic analysis in another *Daphnia* species, *Daphnia galeata,* challenged with a microsporidian parasite, suggests immunity-metabolism trade-offs (Lu et al., 2018).

When it comes to fungal pathogens, the molecular basis of the immune response of *Daphnia* is even more understudied, despite their broad distribution in natural populations and the important impacts they have on host fitness (Ebert, 2005; Shaw et al., 2021; Paraskevopoulou et al., 2022). Among these pathogens, *Metschnikowia bicuspidata* is one of the most prevalent fungal parasites of *Daphnia* (Ebert, 2005; Hall et al., 2010; Hall et al., 2011; Ebert, 2022). This yeast parasite forms needle-shaped ascospores that are horizontally transmitted through ingestion (Ebert et al., 1997; Stewart Merrill & Cáceres, 2018). After ingestion, spores penetrate the gut epithelium and start proliferating via distinct developmental stages in the haemolymph of the animal, ultimately leading to host death (Ebert, 2005; Stewart Merrill et al., 2018). So far, only one transcriptomic study has examined early-stages immune response (within 24 hours of exposure) in *D. dentifera* challenged with *M. bicuspidata*. This study revealed a number of upregulated genes related to cuticle development and defense responses (Terrill Sondag et al., 2023). However, we still lack information whether these immune responses are conserved across *Daphnia* species or how hosts respond beyond early contact with the parasite. Here, we examined gene expression changes across unexposed, exposed-infected and exposed-uninfected individuals to capture differences between immune system activation and metabolic shifts due to infection. Then, we associated these shifts with observed trade-offs in other phenotypic traits, such as host survival and reproduction.

## Material & Methods

### Experimental design

*Daphnia magna* (NO-V-7 clone, Norway) were acclimated under standardized lab conditions (20±1°C, 16:8 L:D) for two generations. The experimental generation was isolated from the third brood of the last acclimated generation and fed daily with *Scenedesmus* sp. *ad libitum* (Izhar & Ben-Ami, 2015). Five days post-birth, 720 juveniles were placed in jars with 20 mL of Artificial *Daphnia* Medium (ADaM; Ebert et al., 1998; Klüttgen et al., 1994) and randomly assigned to one of two treatments: (P) exposed that were inoculated with 1500 spores/mL *bicuspidata* spores for five days, and (C) unexposed that received crushed uninfected *D. magna* as a placebo. Five days post-exposure, all *Daphnia* were transferred to jars filled with 80 mL of fresh ADaM to terminate the exposure period and thereafter medium was replaced three times a week.

### RNA isolation and library preparation

On day 12, infection status was visually determined under a stereomicroscope (Leica DFC 295, Germany) based on *Daphnia’s* external phenotype, with infected individuals developing an opaque color due to the accumulation of fungal spores within the haemolymph. The parasite treatment (P) was further divided into either infected (I) or exposed-uninfected (E) groups. Four biological replicates were selected from each treatment (C, E and I). Each *Daphnia* was homogenized with a sterile plastic pestle in 500 mL of Trizol, and RNA was extracted using a method that combines extraction through Trizol and column precipitation (RNeasy® Mini Kit, QIAGEN), with a double final elution step (20 µL each time). RNA quality and quantity were assessed with Qubit (Thermo Fisher Scientific) and 2200 TapeStation (Agilent), respectively. For library preparation, mRNA was enriched from 300 µL total RNA using the NEBNext® Poly(A) mRNA Magnetic Isolation Module (New England Biolabs), followed by stranded library preparation with the NEBNext® Ultra™ II Directional RNA Library Prep Kit. Libraries were sequenced (single-end, 100 bp) on a NovaSeq6000 Illumina platform at the Crown Genomics Institute, Weizmann Institute of Science (Rehovot, Israel).

### Differential gene expression analysis

We trimmed raw reads with Trimmomatic v0.39 and assessed their quality using Fastqc (Andrews, 2010). Trimmed reads were mapped to the *D. magna* genome (daphmag2.4; GCA_001632505.1) with STAR (Dobin et al., 2013) and gene-level quantification estimates produced by RSEM (Li & Dewey, 2011) (Table S1). Data were imported into R/Bioconductor with the “Tximport” package (Soneson et al., 2015) and gene expression was estimated using the “DESeq2” R package, which performs a size factor normalization to account for differences in sequencing depth among libraries (Love et al., 2014). Genes with zero counts across all samples and those with low counts (< 100) in less than 25% of the samples were filtered out. Before conducting differential expression analysis, we performed a principal component analysis (PCA) to assess sample clustering (Figure S1). We observed that one sample, originally classified as exposed-uninfected (library 383), clustered with infected individuals. Given this discrepancy, we excluded that sample from further analyses, as it was likely an infected individual misclassified due to limitations in infection detection via microscopy. Differential gene expression analysis was performed in three pairwise comparisons: E *vs.* C, I *vs.* C, and I *vs.* E. Significance was based on the Wald test and shrunken log fold-change values with apeglm v1.10.1 (Zhu et al. 2019). Genes were considered differentially expressed (DEGs) at an adjusted p-value < 0.05 and LFC > 0.58. A Venn diagram (Venn.Diagram v1.6.20) was used to depict the overlap of significant DEGs for each pairwise comparison, while their expression profiles were visualized in heatmaps using normalized counts scaled per gene (pheatmap v1.0.12; Kolde, 2018). To identify enriched biological functions in DEGs, we annotated all *D. magna* coding sequences (CDS) using EggNOG (Huerta-Cepas et al., 2019) with an e-value cutoff of 1e−10 (Table S2). Gene Ontology (GO) enrichment analysis was performed with TopGO v2.48.0 (Alexa & Rahnenfuhrer, 2024), using the “classic” algorithm and a minimum genset of three genes. Significantly enriched functions in Biological Process (BP), molecular function (MF), and Cellular component (CC) were inferred using a Fisher’s Exact Test with an adjusted p-value < 0.05. KEGG Orthologs (KO) term enrichment was conducted with clusterProfiler v4.4.4 (Wu et al., 2021) and by applying a p-value < 0.05 cut-off (FDR-adjusted, Benjamini-Hochberg correction).

### Pathway level differential expression analysis

To determine significant changes in pathway-level gene expression, we performed a pathway expression analysis using the “GAGE” v2.46.1 R package (Luo et al., 2009), which identifies differentially regulated pathways by evaluating expression changes across entire gene sets, rather than single genes. The normalized counts from DESeq2 were used and significance was inferred at a False discovery rate (FDR) cutoff < 0.05. Redundancy was reduced by prioritizing pathways with the most distinct and functionally relevant gene contributions using the *esset.grp* function in GAGE package. The R package “Pathview” v3.5.1 (Luo & Brouwer, 2013) was used for visualization.

### Phenotypic data analysis

Survival and reproduction were recorded daily up until the release of the first brood. Thereafter, survival was recorded daily, while reproduction three times per week upon host death. Life-history traits, i.e., age at first reproduction (AFR), brood size at first reproduction (BSFR), offspring production and survival were recorded as proxies of parasite tolerance. AFR was defined as the age of releasing the first brood from the brood pouch. Animals that had died before the first infected individual died were excluded, as this death was likely due to non-natural causes. Animals that were sampled for the RNA-sequencing experiment and animals that died because they were mishandled in the lab were also excluded (Table S3).

Both AFR and Survival were modelled with Kaplan Mayer curves using the *survfit* function in the “survival” v3.3.1 package. Significance was inferred using a Log rank test at a p-value < 0.05 level. BSFR and offspring production were modelled with generalized linear models (GLMs) using a negative binomial distribution and implemented in r using the *glm.nb* function of the “MASS” v7.3.57 package (Venables & Ripley, 2002), with treatment as a fixed factor. Statistical significance was assessed using Type II Wald tests. Post hoc comparisons were computed using the “emmeans” package (Lenth, 2016). All statistical analyses were performed using R v4.2.1, while for visualization, the package ‘ggplot2’ was used (Wickham, 2016).

## Results

### Impact of infection on life-history traits

Exposed-uninfected (E) and unexposed (C) animals lived longer than infected (I) ones (I *vs.* C: p<0.001; I *vs.* E: p<0.001; *Figure 1a*), while there was no difference in survival between those two groups (E *vs.* C: p=0.28; *Figure 1a*). Infected animals reached maturation (AFR) earlier than unexposed (I *vs.* C: p<0.001; *Figure 1b*) and exposed-uninfected (I *vs.* E: p<0.001; *Figure 1b*), while exposed-uninfected matured later than unexposed animals (E *vs.* C: p<0.001; *Figure 1b*). Offspring production (BSFR) was significantly affected by infection, with infected individuals producing less offspring than both exposed-uninfected and unexposed animals (p<0.0001; *Figure 1c*). However, there was no significant difference between unexposed and exposed-uninfected animals (p=0.8352; *Figure 1c*), indicating that exposure without infection does not affect BSFR. Finally, the total number of offspring was also significantly lower in infected animals (I *vs.* C: ratio=23.7, p<0.0001; I *vs.* E: ratio=21.6, p<0.0001; *Figure 1d*). However, there was no significant difference between unexposed and exposed-uninfected (E *vs.* C: ratio=1.1, p=0.6862; *Figure 1d*).

**Figure 1.**
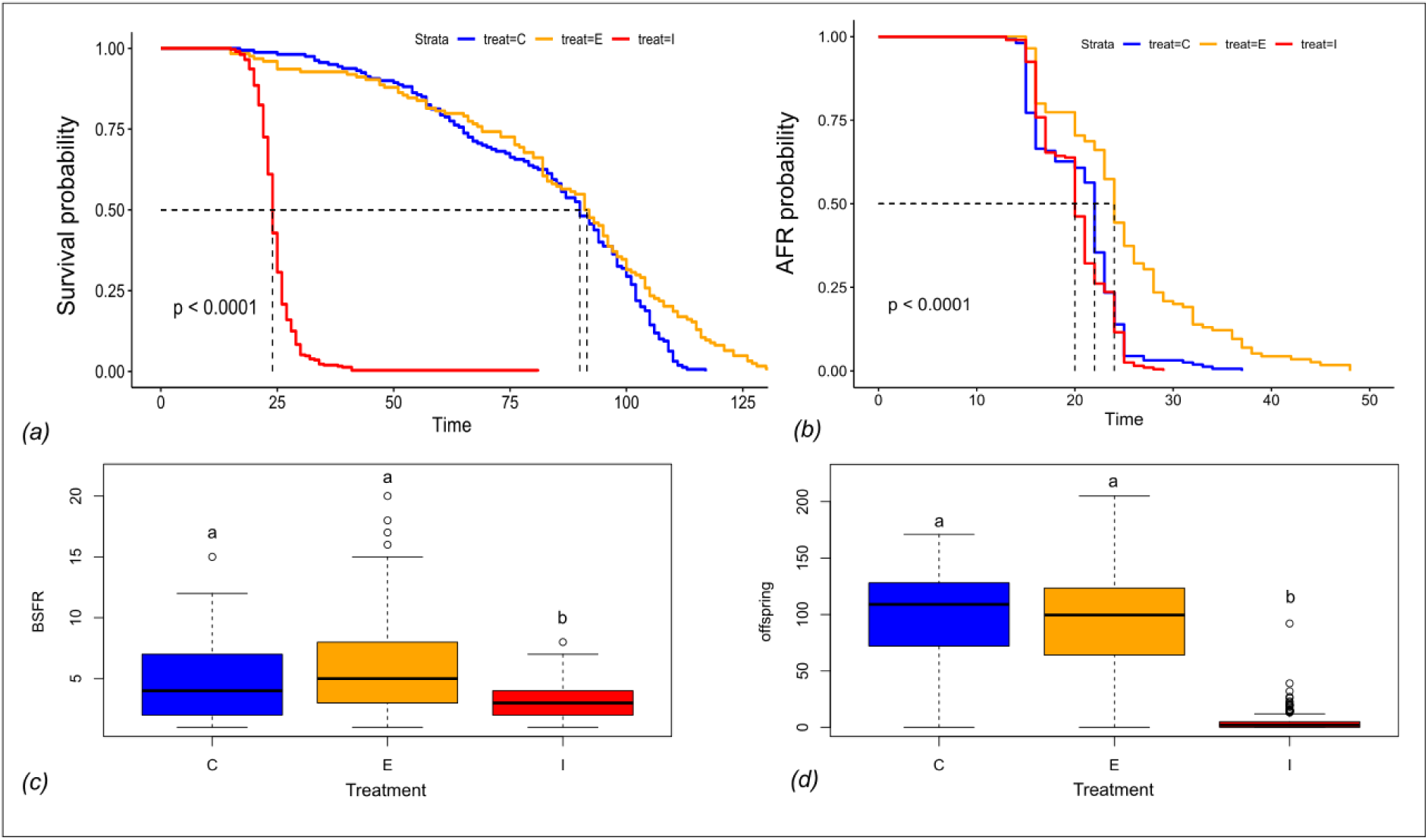
Life-history traits in unexposed (C), exposed-uninfected (E) and infected (I) individuals. *(a)* Kaplan-Meier survival curves for the three groups; the p-value represents the Log-rank test. (*b)* Age at first reproduction (AFR); the p-value represents the Log-rank test. *(c)* Brood size at first reproduction (BSFR); error bars indicate standard deviation (SD). Different letters above groups (*a, b*) indicate statistically significant differences (p < 0.05)*. (d)* Total number of offspring across lifetime; error bars indicate SD. Different letters above groups (*a, b*) indicate statistically significant differences (p < 0.05).

### Impact of infection on gene expression

We identified significant shifts in the transcriptomic landscape associated with parasite exposure and infection (*Figure S2*). In exposed-uninfected individuals compared to unexposed controls (E vs. C), we identified 219 differentially expressed genes (189 upregulated, 29 downregulated; *Figures 2a and S3, Table S4*). Gene Ontology (GO) analysis revealed upregulation of functions related to tissue morphogenesis, molting cycle and cuticle development in exposed-uninfected individuals (*Figure 2b, Table S5, Figure S2a)*. In infected individuals compared to unexposed controls (I vs. C), we identified 760 differentially expressed genes (392 upregulated, 368 downregulated; *Figures 2a and S3, Table S6*). GO analysis revealed upregulation of genes involved in organ morphogenesis, while functions associated with extracellular matrix components and heme biosynthesis were enriched in downregulated genes (*Figure 2b, Table S7*). Seven KEGG terms, including glucose and trehalose transporters (K08145, K14258) and Cytochrome P450 family (K17953, K15001), were enriched in downregulated DEGs (Table S8). In infected individuals compared to exposed-uninfected ones (I vs. E), we captured the most extensive gene expression changes. Specifically, we identified 1,263 differentially expressed genes (586 upregulated, 677 downregulated; *Figures 2a and S3, Table S9*). Seven KO terms were enriched in downregulated genes, including trehalose transporters (K14258), collagen (K19721) and Fibroblast Growth Factor Receptors 1 and 2 (K18496, K18497) (*Table S10*). In contrast, upregulated genes were enriched in 5 KO terms, including cathepsins C and L (K01275, K01365) and Lysosomal Acid Lipase (K12230) (*Table S10)*. GO analysis identified extracellular space-related functions enriched in upregulated DEGs, while genes involved in tissue morphogenesis, epithelial differentiation, chromosome segregation and cell cycle regulation, were downregulated in infected hosts (*Figure 2b, Table S11*).

**Figure 2.**
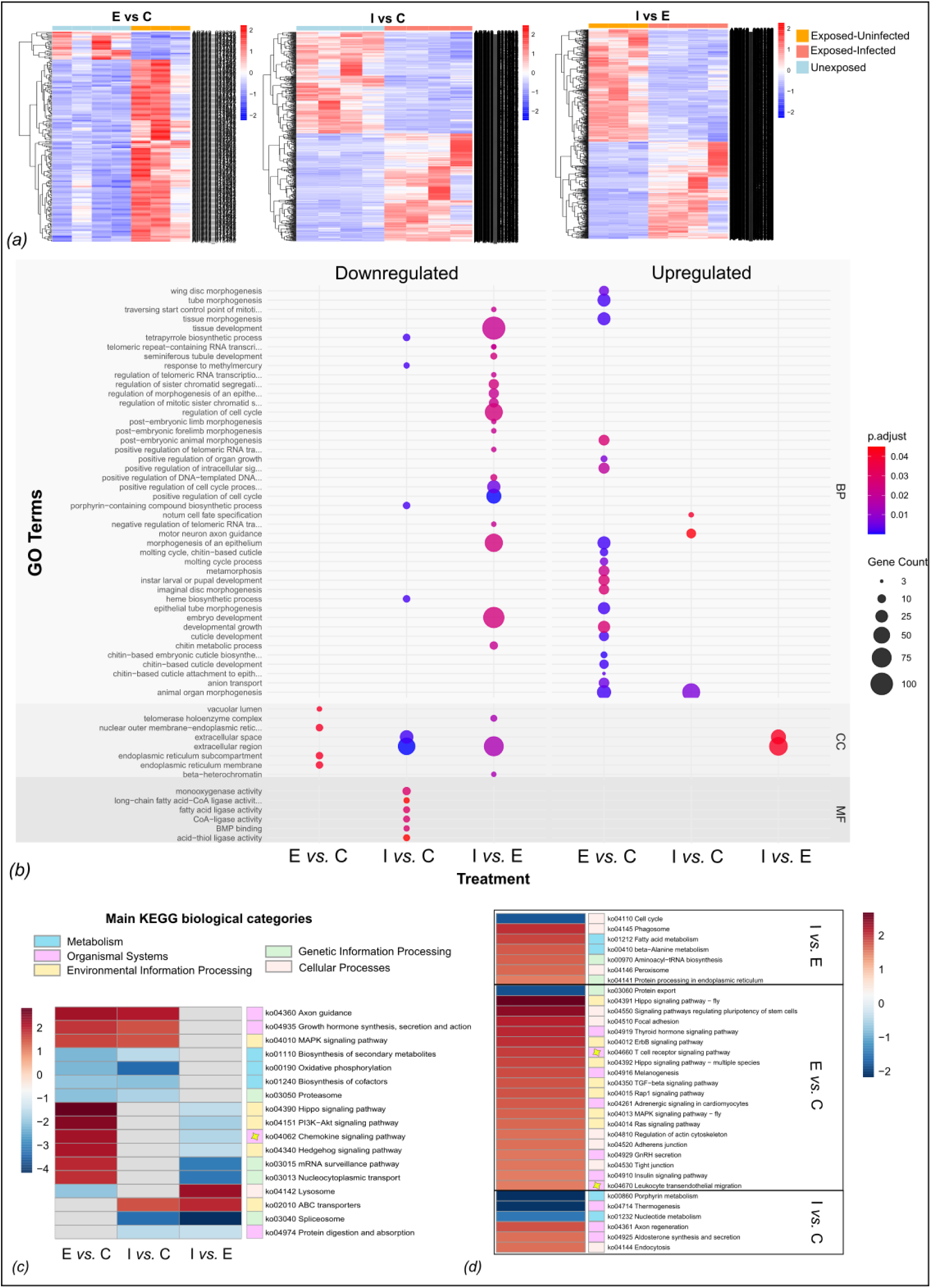
*(a)* Heatmaps showing differentially up- and down-regulated genes across three pairwise comparisons (E vs. C, I vs. C, and I vs. E). For visualization, the normalized to library size gene counts are plotted, with bars representing z-scores. Red indicates upregulation and blue indicates downregulation relative to the first condition in each comparison. *(b)* GO term enrichment analysis across three functional categories: Biological Process (BP), Molecular Function (MF) and Cellular Component (CC). Circle size represents the number of differentially expressed genes (DEGs) associated with each GO term, while color indicates the adjusted p-value. For datasets with more than 20 enriched GO terms, only the 20 most significant ones are shown. *(c)* Heatmap of significantly enriched pathways in two pairwise comparisons. *(d)* Heatmaps of uniquely enriched pathways in each pairwise comparison.

### Impact of infection on whole-pathway expression

Pathway-level expression analysis revealed that exposure to the parasite triggered immune-related responses. Specifically, in exposed-uninfected individuals, we captured an upregulation of pathways annotated to immune system KEGG biological category, such as Leukocyte transendothelial migration (ko04670), T cell receptor signaling pathway (ko04660) and Chemokine signaling pathway (ko04602) (*Figures 2c and 2d*). Similarly, pathways involved in environmental signaling, such as the Hippo, PI3K- Akt and MAPK signaling pathways, together with pathways such as melanogenesis, were upregulated in exposed-uninfected individuals (*Figure 2c*).

In contrast, infection led to distinct metabolic and signaling shifts. ABC transporter pathways and MAPK signaling pathways were upregulated, while pathways related to genetic information processing (e.g., spliceosome, proteasome) as well pathways related to metabolism (e.g., porphyrin metabolism) were downregulated (*Figures 2c and 2d)*. Two pathways were consistently altered across all comparisons (ko03008, ko03010). The Ribosome biogenesis pathway was downregulated in infected hosts, while upregulated in exposed-uninfected individuals. The Ribosome pathway was downregulated in both groups challenged with the parasite (I and E).

## Discussion

### The immune landscape of exposed-uninfected hosts

In exposed-uninfected individuals (E), we detected an upregulation of genes related to tissue morphogenesis, molting and cuticle development. Additionally, the melanogenesis pathway, which regulates melanin biosynthesis, was upregulated. While molting has been suggested as a potential defense mechanism in crustaceans (Shields, 2012; Zuo et al., 2018; Zhang et al., 2021; Xu et al., 2020), it is unclear how molting would benefit the host in the context of *Daphnia-Metschnikowia* interactions, given that the parasite invades through the gut rather than attaching externally. Thus, the upregulation of molting-associated genes is more likely linked to a broader immune response rather than an indication of how frequently *Daphnia* molts. Several studies support this link between molting and immunity in invertebrates. For instance, in crabs, the hepatopancreas—a key metabolic and immune organ—exhibits varying immune activity throughout the molting cycle, suggesting that molting is associated with enhanced resistance to pathogens (Xu et al., 2020; Liu et al., 2022). Additionally, the cuticle serves as a transport and storage site for molecules, such as melanin and prophenoloxidase (PO), which play a key role in invertebrate immunity (Asano & Ashida, 2001; Cerenius & Söderhäll, 2004; Cerenius et al., 2010). In insects, cuticle-related proteins facilitate pathogen encapsulation through melanization, physically restricting their proliferation (Fedorka et al., 2013; Nappi & Christensen, 2005). Additionally, in *Drosophila* and mosquitoes, melanin deposition reinforces the cuticle, thus blocking pathogen penetration (Yassine et al., 2012; Edwards & Eleftherianos, 2023). Finally, in other crustacean (e.g., shrimps), genes related to melanogenesis were upregulated after exposure to fungal and bacterial pathogens. Taken together, these results show that melanogenesis has a conserved role in arthropod immunity (Charoensapsri et al., 2014). Therefore, we argue that the joint upregulation of molting-related genes and melanogenesis may indicate a coordinated immune response, reinforcing epithelial barriers.

Since *Metschnikowia* must penetrate the gut epithelium in order to establish an infection, the observed upregulation of cuticle- and chitin-related genes is more likely associated with gut epithelial defenses rather than exoskeletal molting. Indeed, previous studies have shown that variation in gut epithelium penetrability determines the infection outcome in *Daphnia* (Stewart Merrill et al., 2019; Stewart Merrill et al., 2021a; Stewart Merrill et al., 2021b). In *Drosophila*, gut epithelium renewal is stimulated by bacterial exposure, indicating a direct link between tissue morphogenesis and immunity (Kuraishi et al., 2011; Capo et al., 2019). Since chitin is a major structural component of the gut epithelium, the observed upregulation of chitin-binding and cuticle development GO terms may indicate an increased need for gut epithelial reinforcement—possibly through increased peritrophic matrix turnover.

Our results corroborate with findings in the *D. dentifera*-*M. bicuspidata* system, where cuticle development genes were also upregulated in individuals exposed to the parasite up to 24 h post-exposure (Terrill Sondag et al., 2023). However, our findings contradict previous findings in *Daphnia galeata* and *D. magna*, where no significant upregulation of cuticle-related proteins was observed in response to microsporidia and *P. ramosa* exposure, respectively (Lu et al., 2018; McTaggart et al., 2015). The observed discrepancy among those studies suggests that immune responses involving cuticle and chitin synthesis may be related to how each pathogen infects its *Daphnia* host. For instance, *Metschnikowia* physically penetrates the gut epithelium, while *Pasteuria* and *Caullerya* do not (Ebert, 2005). This highlights the importance of cuticle-associated defenses in invertebrate immunity, particularly in response to pathogens that require epithelial penetration in order to establish infection.

Beyond cuticle-associated responses, parasite exposure triggered an immune system activation in exposed-uninfected individuals. Specifically, we detected enrichment in pathways associated with vertebrate immunity, including Leukocyte transendothelial migration, Chemokine and T-cell receptor (TCR) signaling pathways (Magor & Magor, 2001; Buchmann, 2014; Wang & Knaut, 2014; Shah et al., 2021). In vertebrates, chemokine signaling directs leukocytes toward the endothelium, and leukocyte transendothelial migration enables their passage into tissues. However, invertebrates lack leukocytes. Instead, they employ hemocytes as their primary immune cells, and it is their mobilization that plays a key role in pathogen defense (Lavine & Strand, 2002; Coates et al., 2022). Previous observations in the *Daphnia*-*Metschnikowia* system reported hemocyte aggregation within the haemolymph at sites where *Metschnikowia* spores penetrate the gut (Stewart Merrill et al., 2018). Thus, it is likely that the upregulation of Chemokine signaling and Leukocyte transendothelial migration pathways in exposed-uninfected *Daphnia* constitutes a coordinated hemocyte recruitment response, functionally analogous to vertebrate immune cell migration. A simultaneous activation of PI3K-Akt and MAPK pathways implies that hemocytes were actively mobilized and underwent cytoskeletal rearrangements to migrate and adhere to infection sites. Similarly, TCR signaling is a central component of the vertebrate adaptive immune system, where T cell receptors respond to antigen peptides present on macrophages or to peptides from cytosolic pathogens present on infected cells (Rabb, 2002). While TCRs are unique to vertebrates, some intracellular signaling components involved in T-cell activation (e.g., MAPK, PI3K and NF-κB pathways) are evolutionary conserved across both vertebrates and invertebrates (Li et al., 2011; Gilmore et al., 2012; Martins et al., 2016). The simultaneous upregulation of both MAPK and PI3K pathways in exposed-uninfected hosts suggest that the observed TCR pathway upregulation likely reflects shared components of these signaling pathways rather than any form of immune memory activation in *Daphnia magna*.

Immune activation is energetically costly, which can lead to trade-offs with other fitness-related traits, such as reproduction (Langand et al., 1998; Loker, 2010; Demas & Nelson, 2011). In our study, exposed-uninfected *Daphnia* exhibited delayed reproductive maturation, which suggests that immune activation diverted resources away from reproduction. However, this delay was not translated into long-term reproductive costs, as total offspring production was eventually compensated. Immune-reproduction trade-offs have been observed in other invertebrates, such as molluscs. In snails, an increase in harbored hemocytes during the immune response was associated with reproduction and survival trade-offs (Rigby & Jokela, 2000), while snails resistant to *Exhinosstoma caproni* exhibited delayed maturity compared to susceptible ones (Langand et al., 1998). Recent studies have also shown an energy-related trade-off between immune activation and reproduction, where post-spawning scallops exhibited reduced hemocyte function, likely due to energy depletion following reproduction (Brokordt et al., 2019). This suggests that there is some association between immune activation and reproduction. In our study, the sampling for the RNA-sequencing occurred before the animals first reproduced, therefore, our molecular data most likely captured the energetic investment in immune activation before reproductive maturation. Once this immune investment subsided, reproduction was restored, allowing individuals to compensate early delays. We suggest that immune triggering may temporarily postpone reproduction rather than impose a permanent fitness cost. We also highlight the importance of considering the timing of measurements when assessing immune-related trade-offs, as costs may be transient and stage-specific.

### The immune landscape of infected hosts

While exposed-uninfected individuals showed upregulation of immune-related pathways, infected hosts exhibited major metabolic shifts. Specifically, KEGG Orthology (KO) terms related to glucose and trehalose transporters were enriched in downregulated DEGs in infected *Daphnia*. Glucose and trehalose are sugars that participate in the process of glycolysis in order to produce ATP, the essential energy for cell function. Trehalose is a widely conserved sugar in crustaceans, known for its role in energy metabolism, stress tolerance and immune response (Wang et al., 2016; Huang & Shi, 2023; Jiang et al., 2023; Santos et al., 2024). Trehalose-phosphate synthase (TPS) genes are present in the *Daphnia magna* genome (Huang & Shi, 2023) and disrupting their function impairs the stress response of these animals (Santos et al., 2024). In *Drosophila*, the expression of those transporters increases during hemocyte recruitment, eventually increasing sugar uptake (Kazek et al., 2024). Their downregulation in infected *Daphnia* suggests that infection with *M. bicuspidata* disrupts sugar metabolism. Consequently, the host enters a metabolically compromised state, where an earlier immune activation has exhausted sugar levels in the haemolymph.

Further support for our conjecture is given by the downregulation of genes related to Cytochrome 450, heme biosynthesis and porphyrin metabolism pathway. Porphyrins serve as precursors to heme (Dailey, 1997), which is a crucial component of cytochromes, including Cytochrome P450 enzymes (Fisher et al., 1973) that regulate detoxification, lipid metabolism and immune responses (Nappi & Ottaviani, 2000; Snyder, 2000). The suppression of porphyrin metabolism could directly impair cytochrome-dependent processes, further weakening host defenses and impairing metabolic processes. Upregulation of heme biosynthesis in early infection stages may indicate a host-driven nutritional immunity strategy, where the host limits iron availability to the parasite to prevent its proliferation (Buchmann, 2014; Iatsenko et al., 2020; Núñez et al., 2018). However, once infection is established and host immunity is compromised (as indicated by visual signs of infection in *Daphnia*’s body cavity), heme biosynthesis was significantly downregulated. This, together with the suppression of other functions, such as the ribosomes and ribosome biogenesis, suggests that most likely *Daphnia* suffered from metabolic exhaustion.

This metabolic exhaustion might justify the early reproduction we observed in infected animals. According to the *terminal investment hypothesis*, when organisms face high mortality risk, they increase reproductive allocation to maximize fitness before death (Williams, 1966; Clutton-Brock, 1984). Duffield et al. (2017) propose that this shift is not a fixed response, but rather depends on a dynamic threshold. According to the *dynamic terminal investment threshold* framework, reproductive investment is modulated by both the intensity of the stressor and the energetic state of the host. In our study, downregulation of protein synthesis and energy metabolism genes in infected *Daphnia* showed an energetic exhaustion of the host. This suggests that individuals crossed the threshold for somatic maintenance to be favorable. The simultaneous occurrence of early maturation likely suggests that immune compromise triggered a shift towards immediate reproductive output.

## Conclusions

Our findings showed distinct immune and metabolic responses between exposed-uninfected and infected *Daphnia* hosts, thereby providing insights into the molecular mechanism employed by *D. magna* to combat fungal infections. More precisely, exposed-uninfected individuals activated immune pathways associated with cuticle reinforcement, melanization and hemocyte mobilization, which indicate an immune response that enhances barrier defenses. This immune activation was linked to a delay in reproductive maturation, which was compensated later in life via total reproductive output. Thus, immune triggering in exposed-uninfected *Daphnia* imposed fitness costs that were temporal and transient. In contrast, infected hosts exhibited major metabolic shifts rather than strong immune activation, indicating host metabolic exhaustion that pushed individuals past their dynamic terminal investment threshold. Hence, early reproductive maturation served as a compensatory strategy to maximize fitness in the face of heightened mortality risk.

## Author Contributions

S.P. and S.G, collected the data. S.P analyzed and interpreted the data. S.P. wrote the manuscript with input from S.G. and F.B-A. All authors conceptualized the study. Our study is a lab-based study and all authors are based in the country where the study was conducted.

## Acknowledgements

We want to thank Hila Kobo, Director of the Rosalie and Harold Rae Brown Cancer Research Core Facility at the George S. Wise Faculty of Life Sciences, Tel Aviv University, for helping with the Qubit and Tape Station

## Data Availability Statement

All transcriptomic data have been submitted to the GenBank Sort Read Archive (SRA) under the BioProject number: PRJNA1242537.

