## Supplementary_Figures_S1_S3 for "Transcriptomic landscape reveals immunity related trade-offs in an invertebrate-fungal host-parasite system"

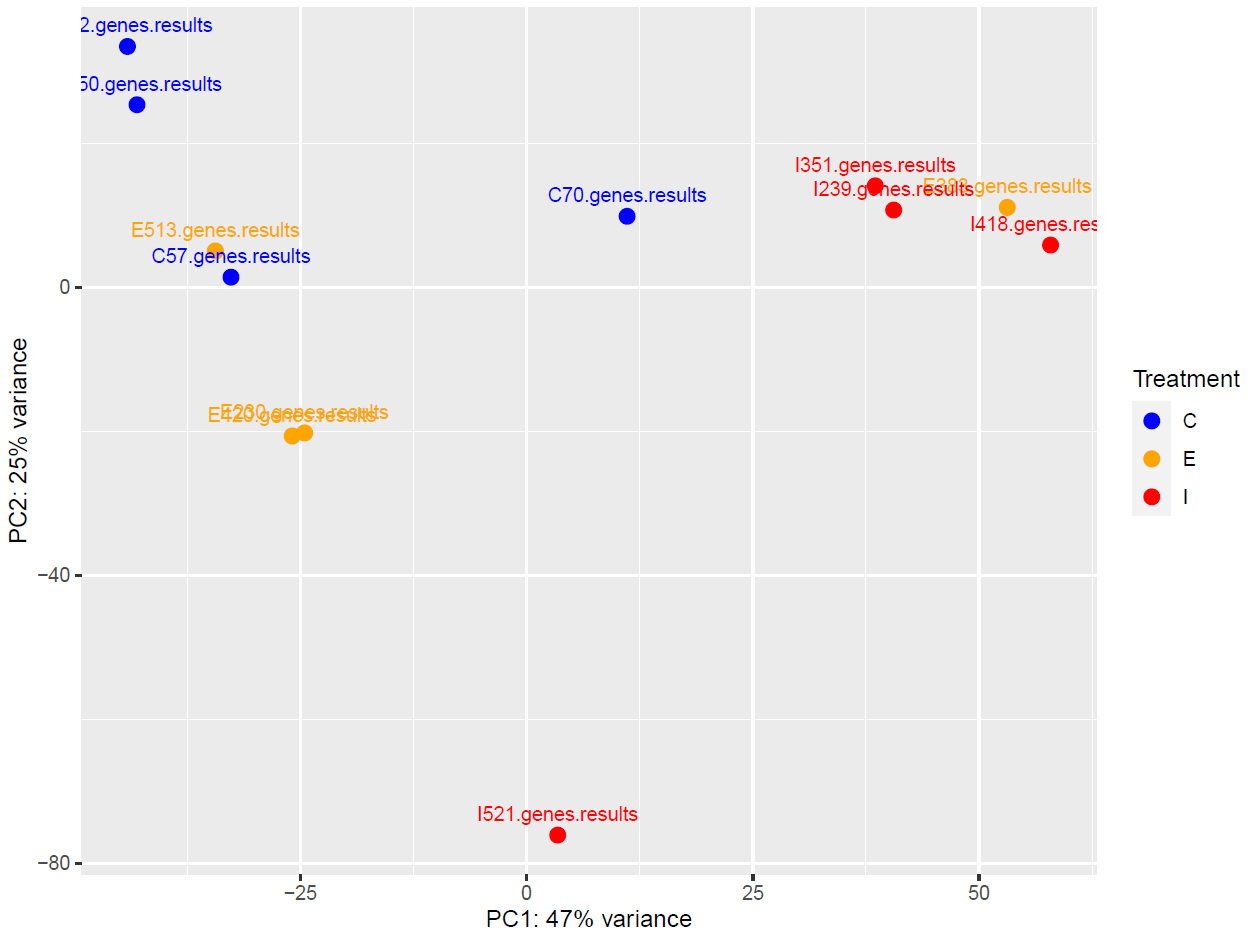

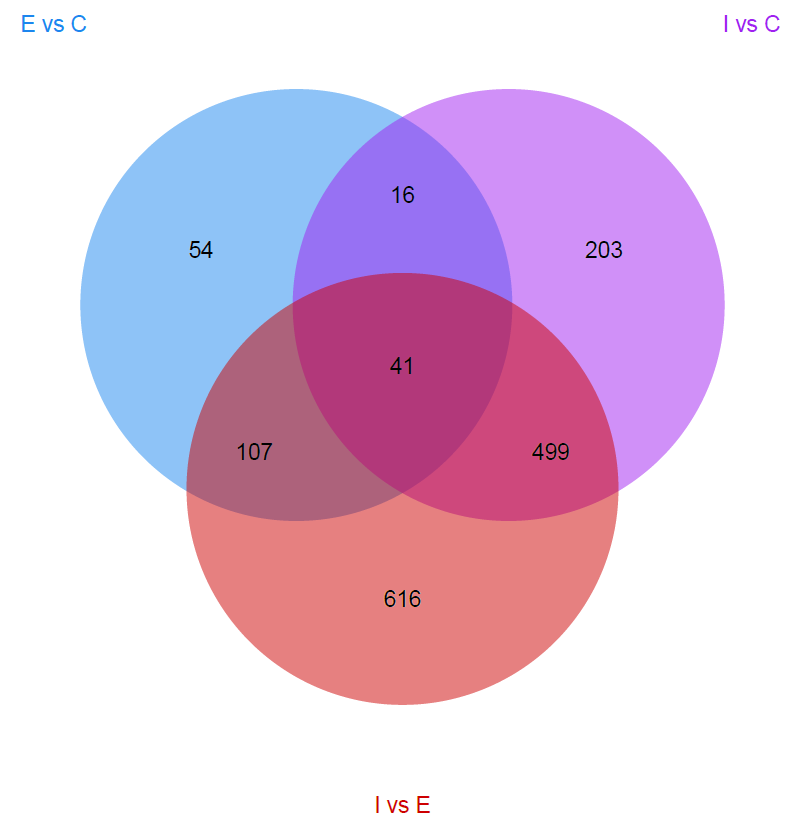
***Figure S1.*** Principal Component Analysis (PCA) showing sample (C, I, E) clustering before we removed sample E_383 from further analysis. Red: infected samples, Orange: exposed-uninfected, Blue: unexposed.

***Figure S2.*** Venn diagram showing shared and unique DEGs among the three pairwise comparisons.

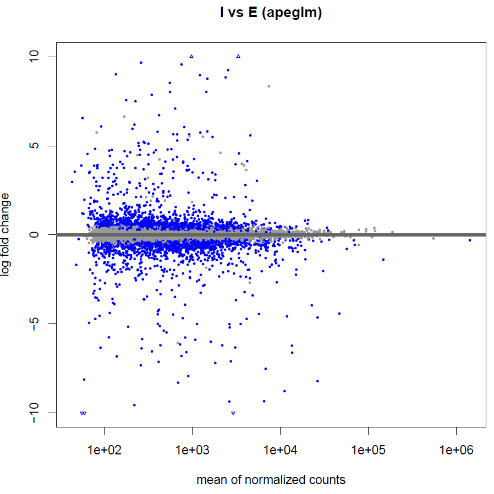

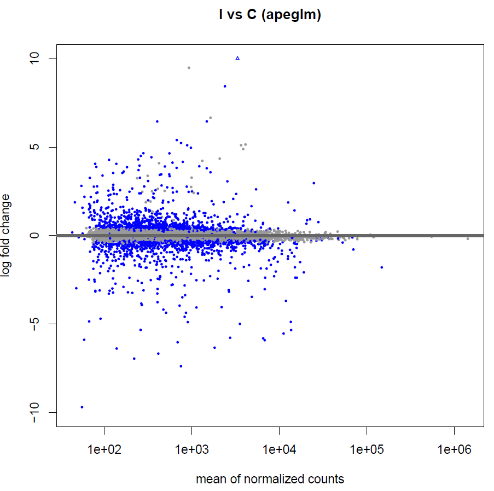

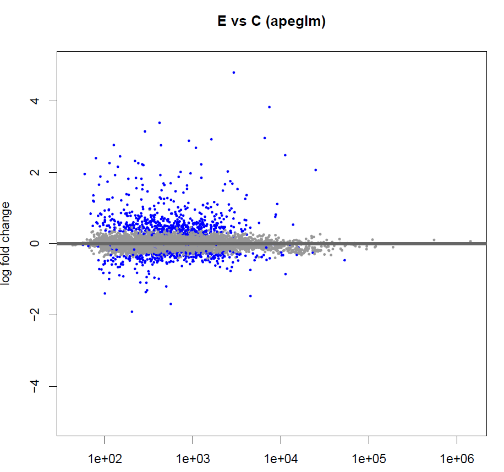

***Figure S3.*** MA plots showing differentially expressed genes (DEGs) across the three pairwise comparisons.
